# Solid tumor CAR T cells engineered with fusion proteins targeting PDL1 for localized IL-12 delivery

**DOI:** 10.1101/2025.04.04.647304

**Authors:** John P. Murad, Lea Christian, Reginaldo Rosa, Yuwei Ren, Eric Hee Jun Lee, Lupita S. Lopez, Anthony K. Park, Jason Yang, Candi Trac, Lauren N. Adkins, Wen-Chung Chang, Catalina Martinez, Carl H. June, Stephen J. Forman, Jun Ishihara, John K. Lee, Lawrence A. Stern, Saul J. Priceman

## Abstract

CAR T cell efficacy in solid tumors is limited due in part to the immunosuppressive TME. To improve anti-tumor responses, we hypothesized that enabling CAR T cells to secrete bifunctional fusion proteins consisting of a cytokine modifier (e.g., TGFβtrap, IL15, or IL12) combined with an immune checkpoint inhibitor (e.g., αPDL1) will provide tumor localized immunomodulation to improve CAR T cell functionality. To that end, we engineered CAR T cells to secrete TGFβtrap, IL15, or IL12 molecules fused to αPDL1 scFv, and assessed in vitro functionality and in vivo safety and efficacy in prostate and ovarian cancer models. CAR T cells engineered with αPDL1-IL12 were superior in safety and efficacy compared to CAR T cells alone and to those engineered with αPDL1 fused with TGFβtrap or IL15. Further, αPDL1-IL12 engineered CAR T cells improved T cell trafficking and tumor infiltration, localized IFNγ production, TME modulation, and anti-tumor responses, with reduced systemic inflammation-associated toxicities. We believe our αPDL1-IL12 engineering strategy presents an opportunity to improve CAR T cell clinical efficacy and safety across multiple solid tumor types.

## Introduction

Chimeric antigen receptor (CAR) T cell therapy for solid tumors has been limited by the suppressive solid tumor microenvironment (TME), which results in poor trafficking of T cells to tumors, T cell exhaustion, inadequate T cell persistence, and restricted engagement of endogenous anti-tumor immune responses ^1–5^. To address these therapeutic challenges, combination strategies with immune checkpoint inhibitors (ICIs; e.g., αPD1, αCTLA4, and αPDL1) and use of immunomodulatory factors (e.g., IL-2, IL-7, IL-12, IL-15, and αTGFβ) have been investigated. ICIs are successful in treating certain malignancies including melanoma, kidney cancers, and non-small cell lung cancers ^6–10^. However, immunologically “cold” cancers, such as metastatic castration-resistant prostate cancer, ovarian cancer, and pancreatic cancer, have shown limited responses to ICIs. Delivery issues, poor biodistribution, and systemic toxicities further complicate the use of ICIs and their combinations in many disease settings. Further, combining CAR T cell therapies with ICIs, which is an obvious combinatorial option, has not produced consistently desirable outcomes ^11–13^. Interleukins play a critical role in both innate and adaptive immune responses, enhancing T and NK cell activity and improving anti-tumor responses in preclinical models ^14–17^. However, clinical applications of many cytokines including IL-12 and IL-15 have been limited due to tumor non-specific activities, raising concerns with cytokine release syndrome (CRS) and other unwanted systemic toxicities ^18–23^. These results highlight the need for unique tumor-restricted approaches to combine these therapies to improve outcomes with solid tumor immunotherapies.

Research by our group and others have developed CAR T cells with integrated ICI and cytokine support for tumor-restricted activity ^24–29^. Our group recently engineered CAR T cells with a membrane-bound form of IL-12 cytokine (mbIL12), increasing local tumor IFNγ and enabling cooperation between cytokine modulation and TME changes ^29^. However, restricted expression of cytokines by adoptively transferred cells may lead to less pronounced TME modifications than with soluble or secreted IL-12. Similar studies have been applied to IL-15 and other cytokines ^16,26,27,30–39^. Approaches like T cell activation-dependent cytokine secretion or anchoring cytokines to immune-related or extracellular matrix proteins showed promise but still may be subject to systemic toxicity concerns or would require exogenous intra-tumoral delivery ^14,18,40,41^. We reasoned that tumor-targeting CAR T cells engineered to secrete bifunctional αPDL1-cytokine fusions would enhance IFNγ induction, subsequently increasing immune checkpoint ligands, such as PDL1, and provide an intra-tumoral ligand for further sequestering cytokine to the local TME to improve the therapeutic index for anti-tumor efficacy.

In the current study, we designed and evaluated CAR T cells that are engineered to secrete bifunctional fusion proteins. We assessed safety and efficacy of this approach using multiple solid tumor CAR T cells engineered with bifunctional proteins comprising a αPDL1 blocking antibody fused with stimulatory molecules (IL12, IL15/Ra, or TGFβ^trap^). These armored CAR T cells exhibit functional PDL1 binding, improved anti-tumor activity, and antigen-specific expansion in tumor co-culture assays. In immunologically “cold” syngeneic prostate and ovarian cancer mouse models ^42–44^, mice receiving CAR T cells engineered with αPDL1-IL12 fusions show superior anti-tumor activity compared with αPDL1-IL15 and αPDL1-TGFβ^trap^. Further, we demonstrate that combination of monotherapies, or CAR T cells secreting non-PDL1 binding controls (αPDL1^mut^-IL12) elicit greater systemic IFNγ and systemic inflammation, while failing to achieve durable anti-tumor responses. In contrast, PDL1 ligation safely facilitated IL-12-mediated increases in intra-tumoral IFNγ, increased CAR T cell presence within the TME, and remodeling of the suppressive TME. We believe our CAR T cell engineering strategy to locally secrete and sequester αPDL1-IL12 fusion proteins within the TME has clinical application across a variety of solid tumor CAR T-cell therapies to safely and dramatically improve therapeutic outcomes.

## Materials & Methods

### Cell lines

The mouse prostate cancer cell line PTEN^-/-^ *Kras* (PTEN-*Kras*) derived from 10-week K-*ras* G12D;PTEN deletion mutant prostate cells was a kind gift from Dr. David Mulholland, The Icahn School of Medicine at Mount Sinai ^42^. The Ras/Myc transformed prostate cancer line, RM9, was a kind gift from Dr. Timothy C. Thompson at MD Anderson Cancer Center and previously described ^45^. ID8, a cell line originated from C57BL/6 mouse ovarian surface epithelial cells was a kind gift from Dr. Karen Aboody, City of Hope National Medical Center. Cell lines were cultured in complete Dulbecco’s Modified Eagles Medium (DMEM) supplemented with 10% fetal bovine serum (FBS, Hyclone), 2 mM L-Glutamine (Fisher Scientific), and 25 mM HEPES (Irvine Scientific) (cDMEM). To generate a more aggressive ID8 tumor line *in vivo*, the parental ID8 cells were engrafted *in vivo* in the peritoneal cavity of mice for approximately 30 days. Unsorted cells harvested from murine ovarian model peritoneal ascites were processed for RBC lysis and analyzed immediately as described. Passaged ID8 cells were then engrafted into tumor naïve mice confirming increased aggressiveness relative to parental.

### DNA constructs, tumor lentiviral transduction, and retrovirus production

Tumor cells were engineered to express firefly luciferase (*ffluc*) by transduction with epHIV7 lentivirus carrying the *ffluc* gene under the control of the EF1α promoter. PTEN-*Kras* cells were transduced to express human PSCA (hPSCA), and ID8 cells were engineered to express tumor associated glycoprotein-72 (TAG72) via transduction with epHIV7 lentivirus carrying the murine *St6galnac-I* gene (mSTn) under the control of the EF1α promoter. mSTn is the unique sialyltransferase responsible for generating surface expression of aberrant glycosylation sialyl-Tn (TAG72) ^46^. The scFv sequence used in the TAG72-CAR construct was obtained from a murine monoclonal antibody clone CC49 that targets TAG72 ^47^. The extracellular spacer domain included the 129-amino acid middle-length CH2-deleted version (ΔCH2) of the murine IgG1 Fc spacer and a murine CD28 transmembrane domain ^47^. The scFv sequence from the murine anti-hPSCA antibody clone (1G8) was used to develop the murine PSCA-CAR construct. The extracellular spacer domain included the murine CD8 hinge region followed by a murine CD8 transmembrane domain ^45^. Both TAG72-CAR and PSCA-CAR had a 4-1BB intracellular co-stimulatory signaling domain. The murine CD3ζ cytolytic domain was previously described ^29,45^. The CAR sequence was separated from a truncated murine CD19 gene (mCD19t) by a T2A ribosomal skip sequence and cloned in a pMYs retrovirus backbone under the control of a hybrid MMLV/MSCV promoter (Cell Biolabs Inc) as previously described ^29,45,47^. Bifunctional fusion proteins or controls contain αPDL1 scFv derived from Avelumab, or mutein non-PDL1 binding variant (αPDL1^mut^) as described previously ^48^. Sequences from full-length murine IL-12(p40p35) ^49^, human IL-15/Ra ^30,48^, and TGFβRII ^30^ were used to construct their respective fusions. Fusions contained linkers using a CH2-deleted version (ΔCH2) of human IgG1 Fc spacer. Retrovirus was produced by transfecting the ecotropic retroviral packaging cell line, PLAT-E (Cell Biolabs Inc.), with addition of murine PSCA-CAR and murine TAG72-CAR retrovirus backbone plasmid DNA using FuGENE HD transfection reagent (Promega). Viral supernatants were collected after 24, 36, and 48 h, pooled, and stored at −80°C in aliquots for future T cell transductions ^45^.

### Murine T cell isolation, transduction, and ex vivo expansion

Splenocytes were obtained by manual digestion of spleens from male heterozygous hPSCA-KI or female C57BL/6j mice. Enrichment of T cells was performed by EasySep mouse T cell isolation kit per manufacturer’s protocol (StemCell Technologies). Single or dual transduction with murine PSCA-CAR, murine TAG72-CAR and/or fusions with αPD-L1, and subsequent expansion were performed as previously described ^29^.

### Flow cytometry

For flow cytometric analysis, cells were resuspended in FACS buffer (Hank’s balanced salt solution without Ca2+, Mg2+, or phenol red (HBSS−/−, Life Technologies) containing 2% FBS and 1x Antibiotic-Antimycotic (AA) (GIBCO; FACS buffer). Single cell suspensions from mouse tissues or tumors were incubated for 20 min on ice with mouse Fc Block (BD 553140). Cells were then incubated with primary antibodies for 30 min at 4°C in the dark with either Brilliant Violet 510 (BV510), Brilliant Violet 570 (BV570), Brilliant Violet 605 (BV605), Brilliant Violet 650 (BV650), fluorescein isothiocyanate (FITC), phycoerythrin (PE), peridinin chlorophyll protein complex (PerCP), PerCP-Cy5.5, PE-Cy7, allophycocyanin (APC), or APC-Cy7 (or APC-eFluor780), eFluor506, PE/Dazzle™ 594, PerCP-eFluor 710, BD Horizon™ Red 718 (R718), Alexa Fluor 488 (AF488), PE-Cy5, -conjugated antibodies. Antibodies against mouse CD3 (BD Biosciences, Cat: 563109, Clone: 17A2), mouse CD4 (Thermofisher, Cat: 340443, Clone: RM4-5), mouse CD8a (BioLegend, Cat: 347313, Clone: 53-6.7), mouse CD19 (BD Biosciences, Cat: 557835, Clone: 1D3), mouse CD45 (BioLegend, Cat: 103145, Clone: 30-F11), mouse CD137 (Thermofisher, Cat: 25-1371-82, Clone: 17B5), mouse NK1.1 (BioLegend, Cat: 108733, Clone: PK163), mouse PD-1 (BioLegend, Cat: 69-9985-80, Clone: J43), mouse LAG3 (BioLegend, Cat: 125227, Clone: C9B7W), mouse TIM-3 (BioLegend, Cat: 119704, Clone: RMT3-23), mouse CD11b (BioLegend, Cat: 101237, Clone: M1/70), CD44 (BD Biosciences, Cat: 103010, Clone: IM7), CD62L (BioLegend, Cat: 104412, Clone: MEL-14), CD80 (BD Biosciences, Cat: 740130, Clone: 16-10A1), mouse I-A/I-E (MHC Class II) (Biolegend, Cat: 64-5321-80, Clone: M5/114.15.2), mouse CD274 (PD-L1) (BioLegend, Cat: 124312, Clone: 10F.9G2), Ly6-C (BioLegend, Cat: 128029, Clone: HK1.4), mouse CD11c (BioLegend, Cat: 117316, Clone: N418), mouse Ly-6G (Biolegend, Cat: 127623, Clone: 1A8), mouse CD103 (BioLegend, Cat: 121426, Clone: 2E7), mouse F4/80 (BioLegend, Cat: 123127, Clone: BM8), mouse IL-12/IL-23 p40 (Thermofisher, Cat: 12-7123-41, Clone:17.8), human IL-15/Ra (R&D Systems, Cat: FAB10900R, Clone: 2639B), human TGFβ Receptor II (BioLegend, Cat: 399703, Clone: W17055E), Purified anti-human TGFβ Receptor II Antibody (BioLegend, Cat: 399702, Clone: W17055E). Cell viability was determined using 4′, 6-diamidino-2-phenylindole (DAPI, Sigma). When necessary, secondary staining of cells was performed by washed twice prior to 30 min incubation at 4°C in the dark. Flow cytometry was performed on a MACSQuant Analyzer 16 (Miltenyi Biotec), and the data was analyzed with FlowJo software (v10.8.).

For intracellular flow cytometry, BD GolgiStop (51-2092KZ) was added to CAR T cells for blocking intracellular protein transport and incubated for 3-4 hours at 37°C. Cells were transferred to a 96-well plate. Reagents and buffers for flow cytometry processing were pre-chilled on ice unless otherwise stated. Cells were washed with FACS buffer and then fixed in 1x BD Cytofix/Cytoperm (51-2090KZ) at 4oC for 20 minutes. Following wash with 1x BD Perm/Wash Buffer (51-2091KZ) twice, cells were stained with intracellular antibody: FITC polyclonal goat anti-human Fc (Jackson ImmunoResearch, Cat: 109-096-008) for 30 min at 4°C. Data was acquired on a MACSQuant Analyzer 16 cytometer (Miltenyi) and analyzed with FlowJo v10.8.

### In vitro PDL1 binding, tumor killing, and functional assays

For PDL1 blocking/binding experiments, PDL1 expression was first induced on RM9 cells for 4 hours via conditioned media collected from CAR T cell:tumor cell co-cultures. PDL1-induced tumors were then co-cultured for 1 hour at room temperature with supernatants collected from supernatants of dual transduced CAR T cells engineered to secrete bifunctional fusion proteins (**Figure 1d**). Levels of PDL1 blockade, from indicated CAR T cell supernatants or relevant positive and negative controls, were measured in a flow cytometry based competitive binding assay using a fluorescently conjugated competitively binding αPDL1 antibody (BioLegend, Clone: 10F.9G2). Simultaneously, PDL1 induced and fusion protein blocked tumors were measured for detection of surface bound fusion cytokines IL15/Ra, IL-12, and TGFβRII. For TGFβRII detection, tumors were first blocked with cold unconjugated anti-TGFβRII antibody to remove tumor receptor background prior to adding CAR T cells supernatants. For tumor cell killing assays, untransduced (UTD), or PSCA-CAR T cells engineered with or without αPDL1, αPDL1-TGFβ^trap^, αPDL1^mut^-IL15, αPDL1-IL15, αPDL1^mut^-IL12, or αPDL1-IL12 were co-cultured at a primary 1:2 effector:tumor (E:T) ratio against PTEN-*Kras* hPSCA mouse prostate tumor cells in complete RPMI without cytokines in 96-well plates (**Figure 1g**). At indicated time points (every 48 hours) co-cultures were analyzed by flow cytometry as described. At these timepoints, replicate plates were rechallenged with an additional 20,000 PTEN*-Kras* hPSCA tumor cells every two days for a total of five tumor challenges. Tumor cell killing by CAR T cells was calculated by comparing CD45-negative DAPI-negative (viable) cell counts relative targets co-cultured with UTD. In addition to flow cytometry measure of co-cultures, cell supernatants were collected from each timepoint to quantify IFNγ by ELISA.

**Figure 1:**
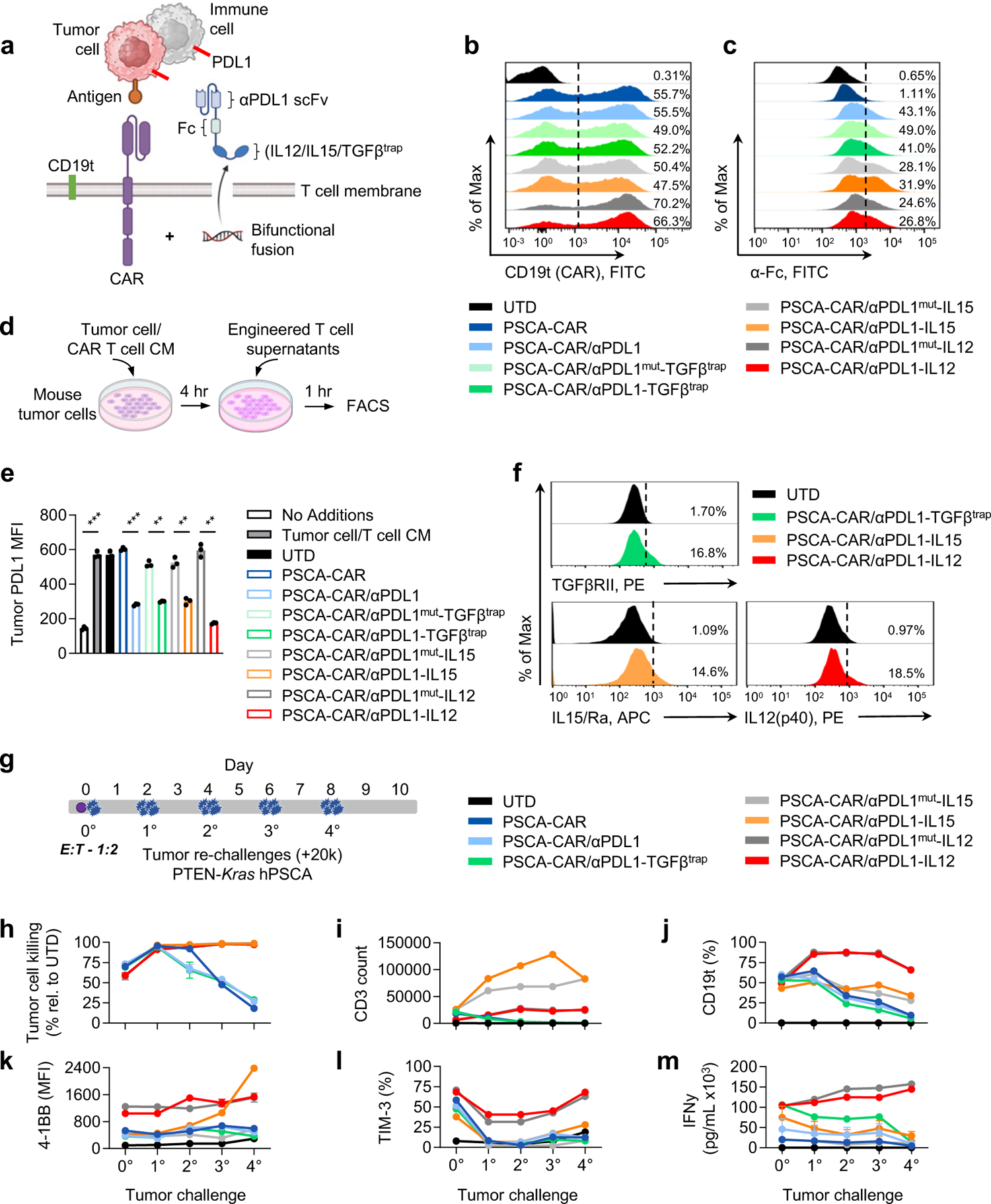
CAR T cells secrete bifunctional fusion proteins and exhibit PDL1 binding and cytokine modulation *in vitro*. **(a)** Illustration of dual transduced CAR T cells engineered to secrete bifunctional fusion proteins with a cytokine modifier (TGFβ^trap^, IL15, or IL12) linked to an αPDL1 targeting scFv. **(b)** Flow cytometry histograms of CAR expression (cell surface CD19t) in indicated untransduced (UTD) or PSCA-CAR conditions. **(c)** Flow cytometry histograms of bifunctional fusion protein expression (intracellular anti-Fc) in indicated conditions. **(d)** Illustration of *in vitro* PDL1 binding/blocking assay on RM9 mouse prostate tumor cells. **(e)** Flow cytometry analysis of mean fluorescent intensity (MFI) of PDL1 on PDL1-induced tumor cells following co-culture with supernatants from indicated UTD, PSCA-CAR, or CAR dual transduced T cells with either αPDL1, PDL1^mut^-TGFβ^trap^, αPDL1-TGFβ^trap^, αPDL1^mut^-IL15, αPDL1-IL15, αPDL1^mut^-IL12, or αPDL1-IL12. **(f)** Flow cytometry analysis of cytokine (TGFβRII, IL15/Ra, or IL12(p40)) co-presentation on the surface of PDL1 induced tumor cells co-cultured with indicated T cell supernatants. **(g)** Illustration of PSCA-CAR T cell and repetitive tumor cell killing assay. UTD, single, or dual transduced PSCA-CAR T cells were co-cultured against antigen-positive PTEN-*Kras* hPSCA cells (E:T = 1:2) followed by flow cytometry analysis and re-challenge with 20,000 tumor cells every 48 hours. **(h)** Flow cytometry analysis of tumor cell killing by CAR T cells engineered with fusions relative to UTD. **(i)** Total CD3+ T cell count, **(j)** percent CD19t+ (CAR+), **(k)** 4-1BB MFI, **(l)** TIM3+, and **(m)** IFNγ by ELISA for each timepoint. n = 3 for each T cell condition and timepoint. Data are presented as mean values ±SEM. Unless otherwise indicated, p-values for pairwise comparisons were generated using an unpaired two-tailed Student’s t-test with assumption of unequal variance where * p<0.05, ** p<0.01, *** p<0.001, **** p<0.0001.

### ELISA cytokine assays

Supernatants from tumor cell killing assays were collected at indicated times and frozen at −80°C for future analysis. Supernatants from all timepoints were thawed and analyzed for murine IFNγ using the murine IFNγ ELISA Ready-SET-GO!^®^ kit (INFγ; Cat no. 88-7314-88, Invitrogen), following manufacturer’s protocol. Mouse serum, plasma, peritoneal ascites, or tumor CD45+ isolated cell supernatant cytokine levels of IFNγ and IL-12 were measured using murine IFNγ and IL-12 ELISA Ready-SET-GO!^®^ ELISA kits (IL12; Cat No. BMS616, Invitrogen), following manufacturer’s protocols. Plates were read at 450 nm using a Cytation3 imaging reader with Gen5 microplate software v3.05 (BioTek). Mouse Luminex® Discovery Assay (15-Plex; Cat no. LXSAMSM-15, R&D Systems) kit was used to evaluate multiple analytes on collected peritoneal ascites from *in vivo* ovarian cancer models. Multiplex cytokine expression data was Log2 transformed and displayed in Balloon plots created using the R package gglot2 version 3.5.1.

### In vivo studies

All animal experiments were performed under protocols approved by the City of Hope Institutional Animal Care and Use Committee. For subcutaneous (s.c.) prostate tumor studies, PTEN-*Kras* hPSCA cells (1.0×10^6^) were prepared in a final volume of 100 uL HBSS^−/−^ and injected under the skin of the abdomen of 6–8 weeks old male heterozygous hPSCA-KI C57BL/6j mice as previously described ^45^. Tumor growth was measured by calipers (length × width × height = mm^3^). For prostate tumor studies, mice were treated by intraperitoneal (i.p.) administration with soluble murine recombinant IL12 (sIL12; 1 μg, PeproTech, Cat: 210-12) once daily for 5 days starting on day of T cell treatment. For *in vivo* i.p. ovarian tumor studies, ID8-mStn (*ffluc* expressing) cells (5.0×10^6^) were prepared in a final volume of 400 uL HBSS^−/−^ and engrafted in >6 weeks old female C57BL/6j (Jackson Laboratories) mice by i.p. injection. Tumor burden was measured via non-invasive bioluminescence imaging (LagoX; Spectral Imaging) and flux signals were analyzed with Aura software (v.4.0, Spectral Imaging). Where indicated, mice were treated i.p. with 100 mg/kg cyclophosphamide (Cy). Mice received intravenous (i.v.) treatment with indicated T cells (1.0×10^6^) in 100 uL final volume 24 hours after Cy pre-conditioning. For ovarian tumor studies, mice were i.p. treated at 14 days post tumor engraftment with indicated T cells (3.0 - 5.0×10^6^) in 400 uL final volume without Cy pre-conditioning. For ovarian tumor studies, mice were co-treated i.p. with CAR T cells and either soluble murine recombinant sIL12 (0.5 μg i.p) or with Avelumab (200 ug i.p.; anti-PDL1, Bavencio, EMD Serono) every other day for total of 3 doses starting on the day of T cell treatment. Mice were euthanized upon reaching s.c. tumor volumes exceeding 1,000 mm^3^ or i.p. tumors showing signs of distress such as a distended belly due to peritoneal ascites, labored or difficulty breathing, apparent weight loss, impaired mobility, or evidence of being moribund.

Peripheral blood was collected from isoflurane-anesthetized mice by retro-orbital (r.o.) bleed through heparinized capillary tubes (Chase Scientific) into polystyrene tubes containing a heparin/PBS solution (1000 units/mL, Sagent Pharmaceuticals). Total volume of each blood draw (approximately 120 uL/mouse) was recorded. Red blood cells (RBCs) were lysed with 1X Red Cell Lysis Buffer (Sigma) according to manufacturer’s protocol and then washed, stained, and analyzed by flow cytometry as described above. When applicable, tumor, liver and spleen were harvested from euthanized mice. Spleen weights were measured for analysis of splenomegaly. Serum from syngeneic prostate and ovarian cancer mouse studies was collected via r.o. bleed in non-heparinized capillary tubes as described above. Blood was kept at room temperature for 30 minutes followed by centrifugation at 6000 x g for 10 minutes at 4°C, aliquoted, and frozen at −80°C until used for serum cytokine ELISA or chemistry analyses. Serum chemistry analysis was performed by running samples on a VETSCAN® VS2 Chemistry Analyzer (Zoetis), using the phenobarbital chemistry panel rotor (Zoetis) for BUN, ALT and AST quantification as described by manufacturer’s protocol. At pre-determined time points or at moribund status, mice were euthanized and tissues and/or peritoneal ascites were harvested and processed for flow cytometry and immunohistochemistry. Peritoneal ascites was centrifuged at 1200 RPM for 10 minutes at 4°C, aliquoted, and frozen at −80°C until used for cytokine ELISA or Mouse Luminex® Discovery Assay multiplex analysis per manufacturer protocol (R&D Systems). For study of immune cells harvested from ovarian i.p. tumor masses, mice from each group were randomly selected and sacrificed at day 6 or 7 post T cell injection. Peritoneal solid tumor masses were physically minced and enzymatically digested via Miltenyi mouse tumor digestion kit and CD45 positive selection was performed using magnetic mouse CD45 MicroBeads as per manufacturer’s protocols (Miltenyi Biotec). Isolated CD45+ cells were analyzed by flow cytometry, and individual replicate supernatants of overnight cultured isolated CD45+ cells from each treatment groups were measured for murine IFNγ secretion via ELISA as described above.

### Immunohistochemistry and Nanostring GeoMx® Digital Spatial Profiling (DSP) analysis

Tumor tissues were fixed for up to 3 days in 4% paraformaldehyde (4% PFA, Boston BioProducts) and stored in 70% ethanol until further processing. Immunohistochemistry was performed by the solid tumor pathology core at City of Hope. Briefly, paraffin-embedded sections (10 μm) were stained with hematoxylin & eosin (H&E, Sigma-Aldrich), CD3 (Ventana, Clone: SGV6), PDL1 (Invitrogen, Clone: JJ08-95), CD4 (Abcam, Clone: EPR19514), CD8 (Cell Signaling, Clone: D4W22), and FOXP3 (Abcam, Clone: EPR22102-37). Images were obtained using the Nanozoomer 2.0HT digital slide scanner and the associated NDP.view2 software (Hamamatzu). For Nanostring GeoMx® DSP, similarly prepared tissues and slides were sent for multiplex protein profiling with spatial context. The Nanostring GeoMx® system provided data center software was used for generating the raw SNR data. Briefly, after QC of the initial data, the segment and target data were filtered followed by generation signal to noise background (SNR) data for downstream analysis. SNR count protein expression data from 12 tumor areas of interest (ROIs) per treatment group were processed and analyzed using R version 4.4.0. The normality of data was assessed using the Shapiro-Wilk test. Protein expression values were z-score transformed and displayed in heatmaps using the ComplexHeatmap R package version 2.20.0 ^50^. ROIs from different treatment groups were clustered using complete-linkage hierarchical clustering on expression data of 39 proteins sub-grouped by cell type or phenotype. The strength and direction of the relationship between the expression of defined pairs of proteins were evaluated using Spearman’s rank correlation method. Unless otherwise specified, a p-value threshold of p < 0.05 was used to determine statistical significance.

### Statistical analysis

In this study, we evaluated CAR T cells for the treatment of solid tumors using *in vitro* T-cell functional assays, as well as syngeneic tumor models in mice. We engineered CAR T cells to secrete bifunctional fusion proetins and evaluated therapeutic efficacy in these model systems. All *in vitro* assays were performed with at least duplicate samples and were repeated in at least two independent experiments. *In vivo* studies were performed using 6- to 8-week-old C57BL/6 or C57BL/6 background huPSCA-KI transgenic mice, using at least three mice per group for all *in vivo* studies to ensure statistical power. Mice were randomized on the basis of tumor volume or bioluminescence imaging to ensure evenly distributed average tumor sizes across each group. *In vivo* experiments were repeated at least twice. For subcutaneous tumor models, survival was based on the maximum tumor size allowed (about 1000mm^3^ in max volume). Data are presented as mean ± standard error mean (SEM), unless otherwise stated. Unless otherwise indicated, p-values for pairwise comparisons were generated using an un-paired two-tailed Student’s t test with assumption of unequal variance where * p<0.05, ** p<0.01, *** p<0.001, **** p<0.0001. GraphPad Prism 10 (GraphPad Software, Inc) was used to generate bar plots and graphs. Data are presented as mean ± standard error mean (SEM), unless otherwise stated. Unless otherwise indicated, p-values for pairwise comparisons were generated using an un-paired two-tailed Student’s t test with assumption of unequal variance where * p<0.05, ** p<0.01, *** p<0.001, **** p<0.0001.

## Results

### CAR T cells secrete bifunctional fusion proteins and exhibit PDL1 binding and cytokine modulation in vitro

Murine splenic T cells were retrovirally-transduced to express murine PSCA-CAR (previously described), CD19t ^45^ and bifunctional fusion proteins (**Figure 1a**). T cell conditions include untransduced (UTD) T cells, PSCA-CAR only, PSCA-CAR/αPDL1, PSCA-CAR/αPDL1^mut^-TGFβ^trap^, PSCA-CAR/αPDL1-TGFβ^trap^, PSCA-CAR/αPDL1^mut^-IL15, PSCA-CAR/αPDL1-IL15, PSCA-CAR/αPDL1^mut^-IL12, or PSCA-CAR/αPDL1-IL12 (**Figure S1a**). All constructs showed comparable CAR T cell transduction as measured by surface expression of mCD19t (**Figure 1b**) ^29,45^, CD8/CD4 phenotypes (**Figure S1b**), and bifunctional fusion proteins and relevant controls as measured by intracellular flow cytometry of the ΔCH2 modified Fc fragment in each fusion protein (**Figure 1c**). Binding of each fusion protein and controls were assessed by an *in vitro* PDL1 binding assay. The murine prostate cancer cell line, RM9, was induced to express PDL1 via conditioned media obtained from tumor cell:CAR T cell co-cultures. Following induction, supernatants from engineered CAR T cells were incubated with tumor cells and assessed by flow cytometry for surface PDL1 binding (**Figure 1d**). Conditioned media resulted in significant induction of PDL1 mean fluorescence intensity (MFI) (**Figure 1e**). Supernatants from CAR T cells engineered with PDL1 fusion proteins, but not αPDL1^mut^ proteins, significantly competed for binding, effectively blocking tumor surface PDL1 and reducing its mean fluorescent intensity (MFI) in our flow cytometry based assay (**Figure 1e**). We also confirmed concurrent surface detection of bound TGFβRII, IL15/Ra, and IL12(p40) via flow cytometry on PDL1-induced tumors (**Figure 1f**).

We next assessed differences in T cell functionality, including tumor cell killing, T cell expansion and activation, and cytokine production by CAR T cells engineered with bifunctional fusion proteins in a repetitive tumor challenge assay *in vitro*. PSCA-CAR T cells engineered with indicated fusion protein or controls were co-cultured with PSCA-positive murine prostate cancer cell line, PTEN-*Kras* hPSCA, at an effector:tumor (E:T) of 1:2. Every 48 hours, co-cultures were analyzed by flow cytometry for tumor cell killing, cell phenotypes, as well as IFNγ secretion by ELISA. Additionally, every 48 hours, T cells were re-challenged with increasing numbers of PTEN-*Kras* hPSCA cells for a total of 4 tumor re-challenges (**Figure 1g**). IL15- and IL12-containing fusions (αPDL1 and αPDL1^mut^) sustained tumor cell killing at nearly 100% (relative to UTD) over each rechallenge as compared to PSCA-CAR only, PSCA-CAR/αPDL1, or PSCA-CAR/αPDL1-TGFβ^trap^ constructs (**Figure 1h**). While IL15 fusions induced the highest total T cell expansion (**Figure 1i**) and CAR T cell expansion each tumor challenge (**Figure S1c**), IL12 fusions induced an enrichment of CAR percentage over time (**Figure 1j**). PSCA-CAR T cells engineered with either αPDL1^mut^-IL12 or αPDL1-IL12 outperformed the remaining constructs over each tumor challenge with higher 4-1BB MFI (**Figure 1k**), and an increase in TIM-3 expression, which is a known downstream target of IL12 signaling (**Figure 1l**) ^51^. Total percentage of 4-1BB, and other markers of exhaustion (PD-1, LAG3), were relatively unchanged (**Figure S1d-f**). While TGFβ^trap^, IL15, and IL12 fusions induced higher initial secretion of IFNγ, only IL12 fusions endowed a sustained increase of IFNγ secretion over the entire tumor challenge assay (**Figure 1m**). The results indicated that CAR T cells engineered to secrete cytokine modifiers IL15 or IL12 enhanced CAR T cell function over CAR T cells alone, but were relatively indistinguishable from their αPDL1^mut^ counterparts, which suggested more extensive *in vivo* assessment was required to determine whether simultaneous PDL1 blockade with the IL12 fusion protein conferred CAR T cell non-autonomous benefits.

### CAR T cells engineered with αPDL1-IL12 fusions exhibit safe and durable anti-tumor responses in a syngeneic murine prostate cancer model

We next assessed the therapeutic efficacy of CAR T cells engineered with various bifunctional fusion proteins *in vivo.* hPSCA-KI mice (previously described) ^45^ were s.c. injected with PTEN-*Kras* hPSCA mouse prostate tumor cells, followed by treatment with Cy (100 mg/kg, i.p.) and indicated CAR T cells (1.0×10^6^, i.v.). Tumor growth was monitored by calipers, and peripheral blood was harvested for flow cytometry and serum cytokine analysis (**Figure 2a**). PSCA-CAR/αPDL1-IL12 T cells showed the greatest anti-tumor response in this model, which was 100% curative in mice treated relative to PSCA-CAR alone or other conditions. Interestingly, PSCA-CAR/αPDL1-IL12 T cells also outperformed PSCA-CAR/αPDL1^mut^-IL12 T cells, which achieved a 50% curative response (**Figure 2b and c**). PSCA-CAR T cells engineered with TGFβ^trap^ and IL15 fusions resulted in little to no therapeutic benefit over CAR T cells alone. Importantly, while mice treated with PSCA-CAR/αPDL1-IL12 T cells showed no significant body weight changes, mice treated with either PSCA-CAR + soluble IL12 (sIL12) or PSCA-CAR/αPDL1^mut^-IL12 T cells exhibited significant weight loss indicating systemic toxicity (**Figure 2d**).

**Figure 2:**
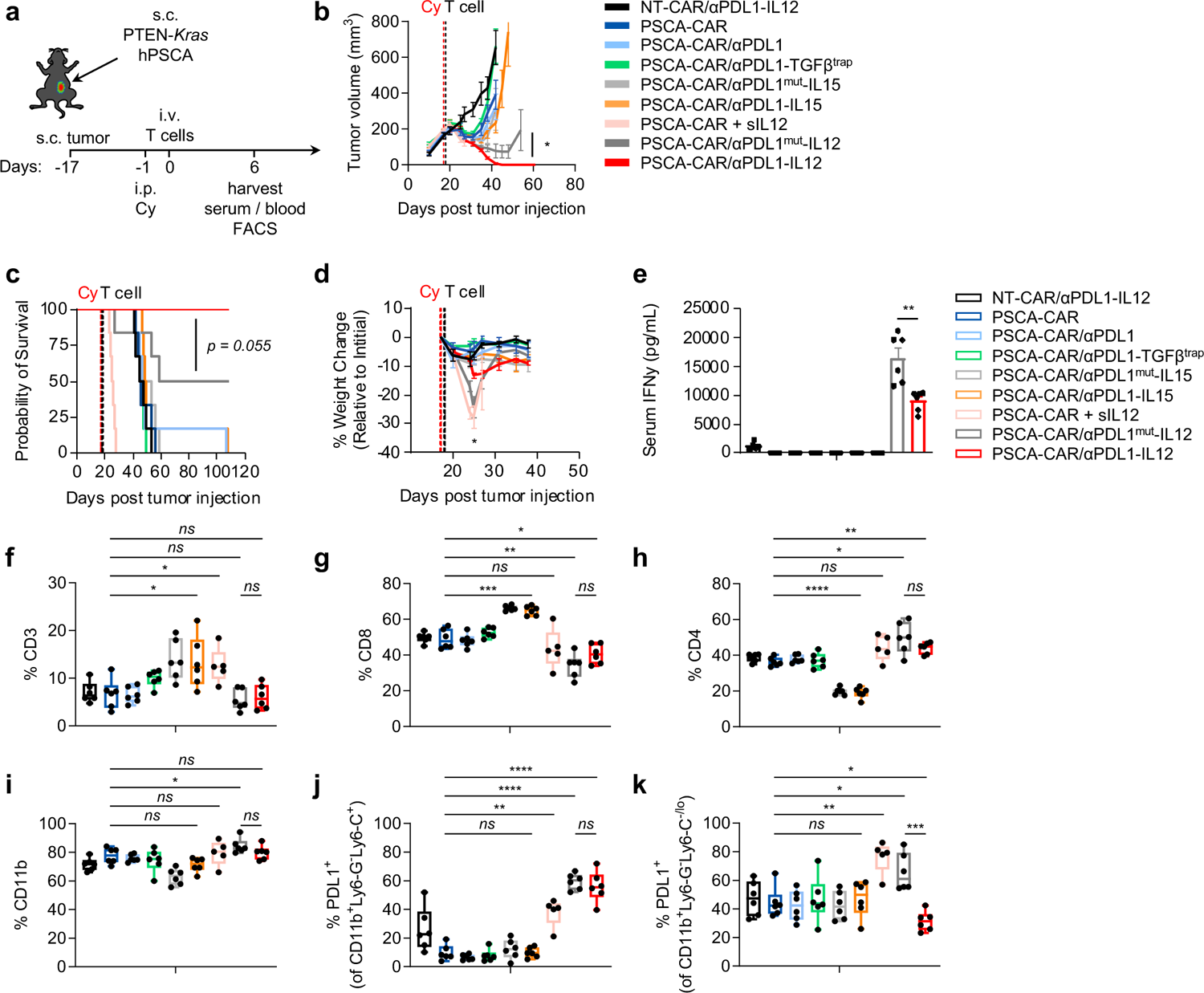
CAR T cells engineered with αPDL1-IL12 fusion exhibit safe and durable anti-tumor responses in a syngeneic murine prostate cancer model. **(a)** Schematic of subcutaneous (s.c.) PTEN-*Kras* hPSCA tumor engraftment, Cy pre-conditioning (100mg/kg, i.p.), and intravenous (i.v.) treatment with non-targeting CAR (NT), PSCA-CAR alone, or PSCA-CAR engineered with αPDL1, αPDL1-TGFβ^trap^, αPDL1^mut^-IL15, αPDL1-IL15, αPDL1^mut^-IL12, or αPDL1-IL12 (1.0×10^6^ CAR+ cells per condition). **(b)** Average tumor volumes (mm^3^) at indicated days post tumor injection for treatment groups. p-value at day 35 post tumor injection comparing PSCA-CAR/αPDL1^mut^-IL12 and PSCA-CAR/αPDL1-IL12. **(c)** Kaplan-Meier survival plot for mice in each indicated group. n ≥ 6 mice per group, p-value comparing TAG72-CAR/αPDL1^mut^-IL12 and TAG72-CAR/αPDL1-IL12 using a Log-rank (Mantel-Cox) test. **(d)** Percent body weight change in indicated groups relative to pre-treatment weight. **(e)** Serum IFNγ by ELISA on day 6 post T cell treatment. **(f-j)** Flow cytometry analysis of peripheral blood at 6 days post T cell treatment. **(f)** Percent of CD3+ T cells, **(g, h)** percent of CD8+ and CD4+ T cells, **(i)** percent of CD11b+ myeloid cells, **(j-k)** percent PDL1-positive cells gated on **(j)** CD11b^+^Ly6G^-^Ly6C^+^ and **(k)** CD11b^+^Ly6G^-^Ly6C^-/lo^ monocyte subsets. Unless otherwise indicated, p-values for pairwise comparisons were generated using an un-paired two-tailed Student’s t test with assumption of unequal variance where * p<0.05, ** p<0.01, *** p<0.001, **** p<0.0001.

Mice treated with PSCA-CAR/αPDL1^mut^-IL12 T cells had significantly higher systemic IFNγ levels relative to PSCA-CAR/αPDL1-IL12 (**Figure 2e**). These findings suggest that PDL1 binding in the TME resulted in reduced systemic IL-12 effects. Serum analysis of AST and ALT showed modest changes across treatment groups, but a trend of increased BUN, which may suggest mild impairment in kidney function in PSCA-CAR/αPDL1^mut^-IL12 T cells and PSCA-CAR T cells + sIL12-treated mice (**Figure S2a-c**). Flow cytometric analysis of blood lymphoid populations showed CAR T cells engineered with IL15 fusions resulted in significantly higher frequencies of circulating CD8 T cells and 41BB expression (**Figure 2f-g and Figure S2d**) but without improved anti-tumor response or survival. Mice treated with PSCA-CAR T cells and sIL12 or engineered with either αPDL1^mut^-IL12 or αPDL1-IL12 showed a significant increase of peripheral blood CD4 T cells (**Figure 2h**). For all indicated treatments, circulating T cells were not CAR positive (**Figure S2e**), as we have observed previously ^45^. For non-CAR CD8^+^ T cells in circulation, we saw minor increase in PD-1 expression in αPDL1^mut^-IL12 and αPDL1-IL12 treatment, significant increases in TIM-3 expression for sIL12, αPDL1^mut^-IL12, and αPDL1-IL12, and slightly lower LAG3 expression relative to PSCA-CAR T cell treatment alone (**Figure S2f-h**). Analysis of circulating myeloid cells in mice showed modest changes in total CD11b^+^ cells in all treatment groups (**Figure 2i**). Mice treated with PSCA-CAR/αPDL1-IL12 T cells showed significant reduction in PDL1 expression among the circulating Ly6C^-/lo^ monocytic myeloid cells (**Figure 2k and Figure S2i-k**), which likely represent competitive fusion protein binding on tumor-experienced and -egressed cells ^52,53^. These results suggest that PSCA-CAR T cells engineered with αPDL1-IL12 exhibit distinct local tumor delivery and subsequent binding to PDL1-positive cells.

### TAG72-CAR T cells engineered with αPDL1-IL12 promote durable anti-tumor responses and exhibit tumor-restricted effects in a syngeneic murine ovarian cancer model

To expand our *in vivo* therapeutic efficacy studies, we evaluated TAG72-CAR T cells engineered with bifunctional fusion proteins in our established ID8 ovarian cancer peritoneal metastasis model (**Figure 3a**). Mice were i.p. injected with an aggressive, *ex-vivo* passaged (**Figure S3**), TAG72-positive ID8 murine ovarian cancer cell line expressing firefly luciferase (ID8-mSTn/ffluc). At 14 days following injection, mice were treated i.p. with either engineered non-targeting (NT)-CAR/αPDL1-IL12 T cells, TAG72-CAR T cells alone, or TAG72-CAR T cells engineered with αPDL1^mut^-IL12 or αPDL1-IL12. Non-invasive imaging of tumors showed TAG72-CAR/αPDL1-IL12 had a dramatic and sustained anti-tumor response (**Figure 3b-d**) resulting in 100% survival rate beyond 150 days post tumor injection (**Figure 3e**). In contrast to αPDL1-IL12, mice treated with TAG72-CAR/αPDL1^mut^-IL12 showed heterogenous anti-tumor responses and reduced overall survival (**Figure 3e**). Mouse weight changes were minimal among all treatment groups (**Figure 3f**), likely due to the regional delivery of CAR T cells compared with systemic delivery as well as the regional tumor development versus local disease in the prostate model (Figure 2).

**Figure 3.**
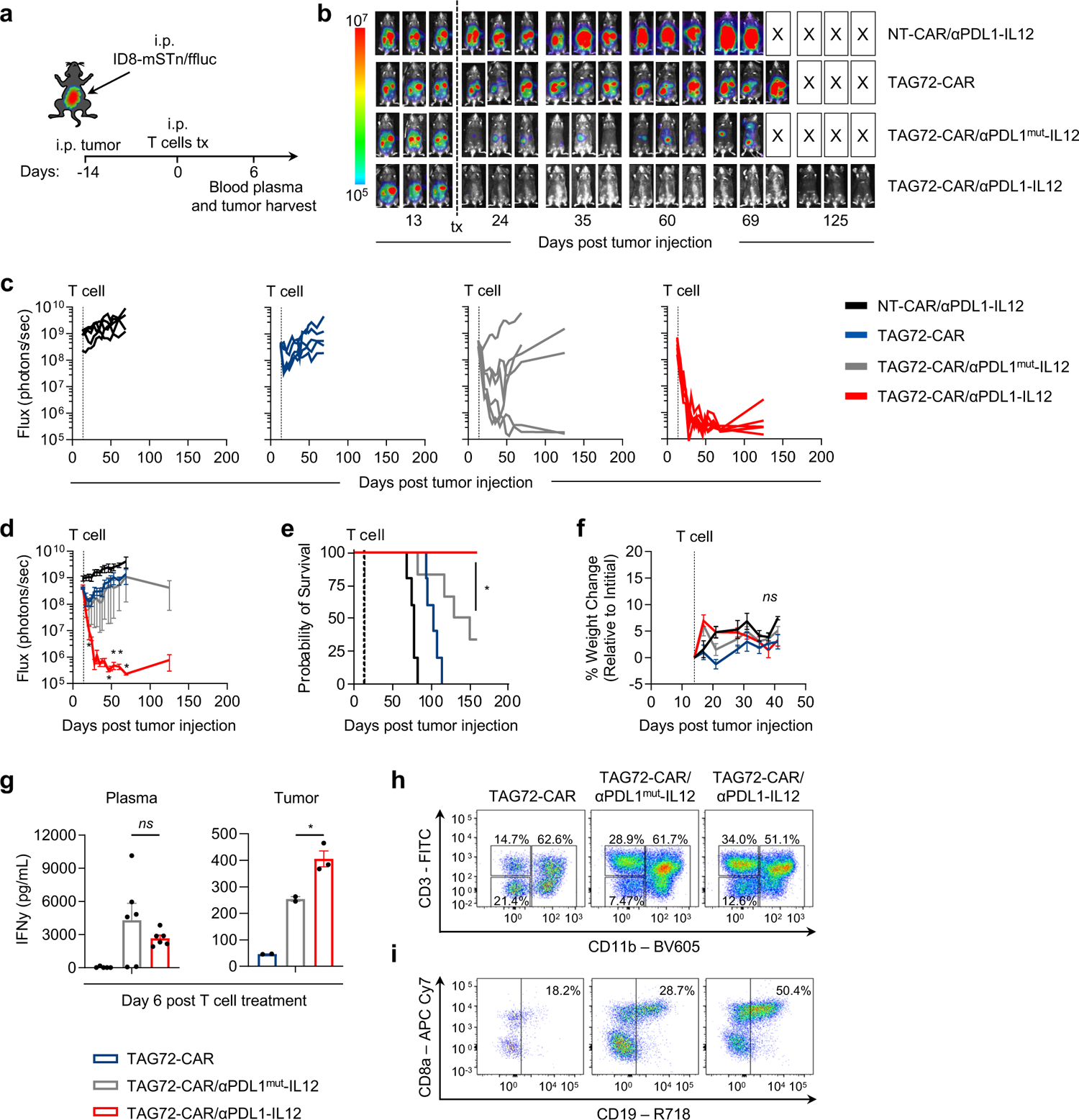
TAG72-CAR T cells engineered with αPDL1-IL12 promote durable anti-tumor responses and exhibit tumor-restricted effects in a syngeneic murine ovarian cancer model. **(a)** Schematic of intraperitoneal (i.p.) ID8-mSTn/ffluc tumor model treatment and harvest timepoints. **(b)** Representative flux images of tumor-bearing mice treated with NT-CAR/αPDL1-IL12, TAG72-CAR, TAG72-CAR/αPDL1^mut^-IL12, TAG72-CAR/αPDL1-IL12 (5.0×10^6^ CAR+ cells; i.p.) as indicated. **(c)** Quantification of flux (photons/sec) from each mouse per group. **(d)** Average flux. p-value at indicated timepoints comparing TAG72-CAR/αPDL1^mut^-IL12 and TAG72-CAR/αPDL1-IL12. **(e)** Kaplan-Meier survival plot. n ≥ 6 mice per group, p-value comparing TAG72-CAR/αPDL1^mut^-IL12 and TAG72-CAR/αPDL1-IL12 using a Log-rank (Mantel-Cox) test. **(f)** Percent body weight change in each treatment group relative to pre-treatment weight (ns; not significant differences between treatment groups at all time). **(g)** IFNγ by ELISA of mouse plasma (n = 3-6 per group; left panel) and intra-tumoral CD45+ isolated cells (n = 3 per group; right panel) at day 7 post T cell injection. Representative flow cytometry plots of **(h)** percent of CD3+ and CD11b+ and **(i)** percent of CD8+ and CD19t+ (CAR+) T cells within intra-tumoral CD45+ isolated cells from indicated groups at day 7 post T cell injection. All data are presented as mean ± SEM. Unless otherwise indicated, p-values for pairwise comparisons were generated using an un-paired two-tailed Student’s t test with assumption of unequal variance where * p<0.05, ** p<0.01, *** p<0.001, **** p<0.0001.

Analysis of blood plasma collected from mice showed a trend of higher systemic inflammation with higher IFNγ in TAG72-CAR/αPDL1^mut^-IL12 relative to TAG72-CAR/αPDL1-IL12 T cell treatment (**Figure 3g; left panel**) similar to our *in vivo* prostate model observations. In a selection of mice harvested at day 6 post treatment, solid tumor masses found within the upper omental region of the peritoneum were digested to single cells and magnetic bead isolated for CD45-positive immune cells. Local IFNγ secretion from tumor isolated CD45-positive cell supernatants, in *ex vivo* culture, were significantly higher in mice treated with TAG72-CAR/αPDL1-IL12 T cells in contrast to TAG72-CAR/αPDL1^mut^-IL12 or TAG72-CAR T cells alone (**Figure 3g; right panel**). Representative flow cytometry analysis of the same isolated CD45 cells showed similarly increased CD3^+^ T cells in TAG72-CAR/αPDL1^mut^-IL12 and TAG72-CAR/αPDL1-IL12 groups (**Figure 3h**), with the highest CAR T cell percentages observed in the TAG72-CAR/αPDL1-IL12 group. Analysis of immune cell populations isolated from the peritoneal ascites from mice harvested at day 12 post T cell treatment in mice treated at day 35 post tumor injection (**Figure S4a**) showed similar trends in IL12-related increased T cell frequencies, but interestingly unlike within the tumor itself, there was a non-significant presence of CAR^+^ T cells (**Figure S4b-d**). Within the ascites of TAG72-CAR/αPDL1-IL12 T cell treated mice, total myeloid (CD11b^+^) cells were relatively unchanged but exhibited decreased PDL1 MFI (**Figure S4e and f**) as a result of fusion protein αPDL1 competitive binding blockade. Further, a majority of these Ly6G^-^ Ly6C^+^ myeloid cells are F4/80^-/lo^ which also exhibit decrease in PDL1 expression relative to αPDL1^mut^-IL12, and this trend is similarly found in F4/80^high^ subsets (**Figure S4g-j**). These results highlight sustained local tumor IFNγ production with αPDL1-IL12 engineered CAR T cells, which in turn enhances tumor infiltration of CAR and non-CAR T cells contributing to the durable anti-tumor responses observed in this model.

### CAR T cells engineered with αPDL1-IL12 fusion elicit distinct changes in the TME

To assess TME changes following CAR T cell treatment, we employed immunohistochemistry (IHC) and Nanostring GeoMx ® spatial proteomic analysis on tumors collected 7 days after T cell treatment in the syngeneic ovarian cancer model as previously described. Tumors from TAG72-CAR/αPDL1^mut^-IL12 and TAG72-CAR/αPDL1-IL12 T cell groups showed increased intra-tumoral CD3, CD4, and CD8 T cells relative to TAG72-CAR T cells alone, with low total frequencies of Foxp3^+^ regulatory T cells relative to total immune cells (**Figure S5**). Furthermore, tumor PDL1 was markedly increased in αPDL1^mut^-IL12 and αPDL1-IL12 groups as a likely consequence of increased local IFNγ. To further interrogate changes in the TME, Nanostring GeoMx ® spatial proteomic analysis was performed. A total of 12 tumor specific regions of interest (ROI) from three replicate mice per group were selected (**Figure 4a and Figure S6**). ROI values were z-score transformed from signal-to-noise ratio adjusted (SNR) counts of detected proteins. A heatmap subdivided into major cell types, phenotype, or status was then generated and unsupervised clustering was applied to organize treatment groups. In mice treated with TAG72-CAR/αPDL1-IL12 and TAG72-CAR/αPDL1^mut^-IL12 T cells, unbiased clustering tightly associated with treatment type (red treatment group and dark gray treatment ROI’s respectively), indicating consistent effects per treatment and per ROI (**Figure 4b**).

**Figure 4:**
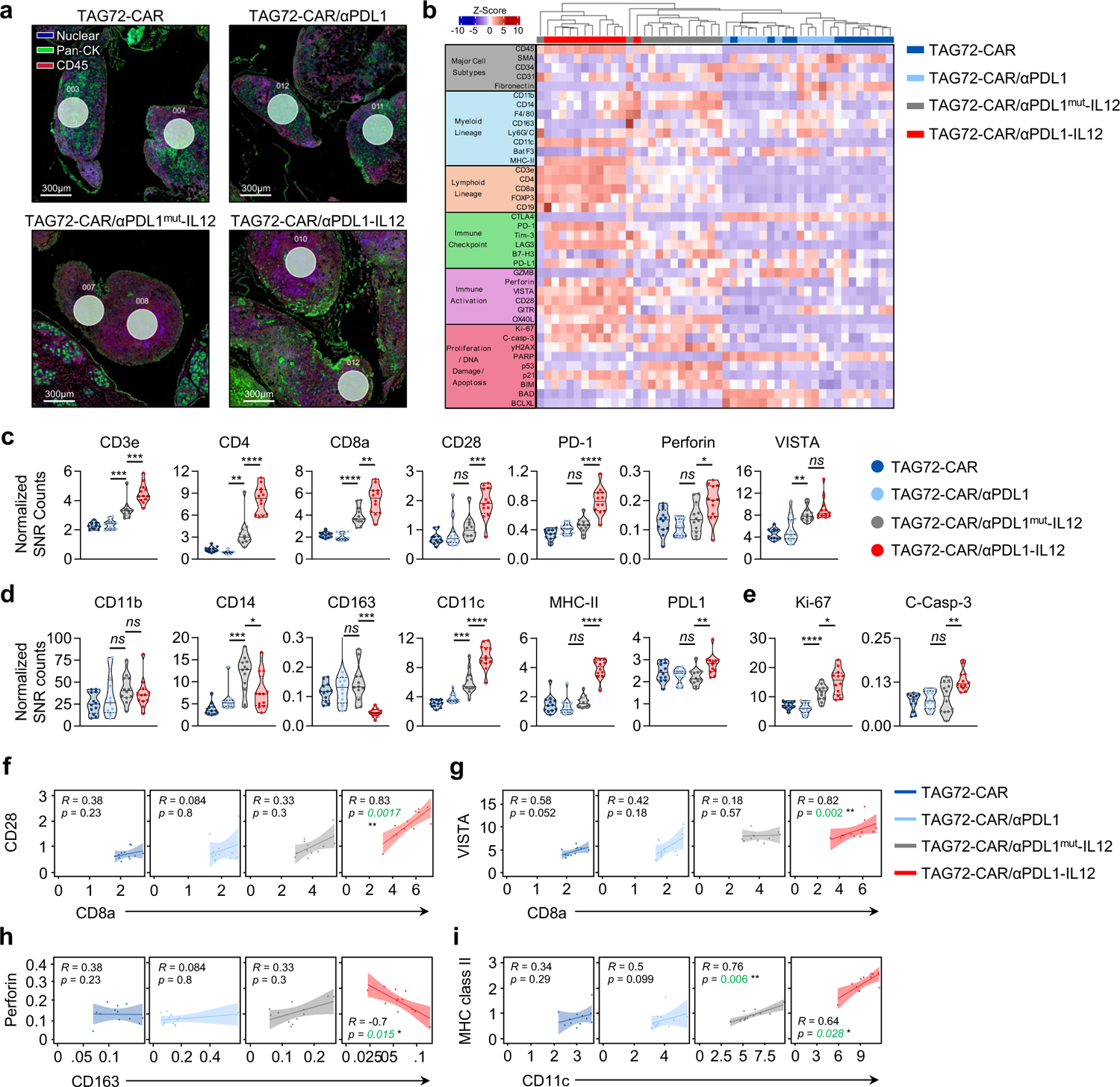
CAR T cells engineered with αPDL1-IL12 fusion elicit distinct changes in the TME. **(a)** Representative Nanostring GeoMx® captured immunofluorescence images and representative regions of interest (ROI’s) from tumor sites in tissues harvested at day 7 post T cell treatment from TAG72-CAR, TAG72-CAR/αPDL1, TAG72-CAR/αPDL1^mut^-IL12, and TAG72-CAR/αPDL1-IL12 treatment groups. Nuclear stain (blue), Pan-CK (green), CD45 (red). Scale bars represent 300 µm. **(b)** Heatmap of z-score transformed protein quantification generated from normalized signal-to-noise ratio corrected counts (SNR counts) within each group (12 ROI’s per treatment group). ROIs from treatment groups were clustered based on the expression of proteins sub-grouped by cell type or phenotype (left panel). Violin plots of normalized SNR counts for individual proteins quantified per ROI per treatment group are shown for **(c)** CD3e, CD4, CD8a, CD28, PD-1, Perforin, VISTA, **(d)** CD11b, CD14, CD163, CD11c, MHC Class II, and PDL1 and **(e)** Ki-67 and Cleaved-Caspase-3 (C-Casp-3). **(f-i)** Spearman correlation plots generated from normalized SNR counts indicating strength of either positive or negative correlations (Spearman’s correlation coefficient, denoted as R) and statical significance of selected correlations per ROI of **(f)** CD28 and CD8a, **(g)** VISTA and CD8a, **(h)** Perforin and CD163, and **(i)** MHC Class II and CD11c. n = 12 ROI’s per treatment group representing n = 3 treated mice per group. Unless otherwise indicated, p-values for pairwise comparisons were generated using an un-paired two-tailed Student’s t test with assumption of unequal variance where * p<0.05, ** p<0.01, *** p<0.001, **** p<0.0001.

In the TAG72-CAR/αPDL1-IL12 T cell treatment group, there was a marked enrichment of proteins related to lymphoid lineage, immune checkpoint, and T cell activation status. This increase was less prominent in TAG72-CAR/αPDL1^mut^-IL12 T cell, TAG72-CAR/αPDL1 T cell, or TAG72-CAR T cell only groups (**Figure 4b**). Moreover, TAG72-CAR/αPDL1-IL12 T cell treatment resulted in significantly greater intra-tumoral lymphoid cell and activation status via measure of CD3e, CD4, CD8a, CD28, PD-1, perforin, VISTA, and Ki-67 in comparison to TAG72-CAR/αPDL1^mut^-IL12 (**Figure 4c**). All treatment groups had relatively similar counts in total CD11b, however, we found that TAG72-CAR/αPDL1-IL12 showed significantly reduced counts in myeloid CD14 positivity ^54,55^, but higher dendritic cells as measured by CD11c (**Figure 4d**). Further, TAG72-CAR/αPDL1-IL12 T cell groups showed significantly lower counts of suppressive M2-like marker CD163 and significantly higher counts of MHC class II (**Figure 4d**) ^54–56^. Of note, only the TAG72-CAR/αPDL1-IL12 T cell treatment group induced a significant increase in measured total PDL1 protein (intracellular and extracellular) within the tumor ROI’s selected, supporting the higher observed levels of local IFNγ. Lastly, total proliferative marker Ki-67 and cell apoptotic marker cleaved caspase-3 was increased in TAG72-CAR/αPDL1-IL12 (**Figure 4e**).

To gain a better understanding of co-expression patterns within tumor tissues for all treatment groups, a spearman correlation analysis was performed. Statistically significant and strong positive or negative correlations were assessed and imply stronger intra-tumoral T cell activation responses in the TAG72-CAR/αPDL1-IL12 T cell group. We observe CD8a T cell presence strongly correlated with both T cell co-stimulatory activation marker CD28 (spearman R = 0.83, p = 0.0017) and T cell V-domain immunoglobulin suppressor of T cell activation (VISTA; R = 0.82, p = 0.002) relative to all other treatment groups (**Figure 4f and g**). Furthermore, in the TAG72-CAR/αPDL1-IL12 T cell group, we observed a negative correlation between greater perforin expression (commonly associated with T cell killing) and lower M2-like suppressive myeloid CD163 expression (**Figure 4h**; R = −0.7, p = 0.015) ^56^. We also observed a strong correlation of MHC class II expression with CD11c in the TAG72-CAR/αPDL1-IL12 T cell group compared to the other groups (**Figure 4i**). Together, these data support a cell extrinsic mechanism of local TME modulation by CAR T cells engineered with αPDL1/IL12 fusions, resulting in improved immune T cell function, shifts in myeloid cell status, and other immune correlates.

### Improved safety of CAR T cells engineered with αPDL1-IL12 compared with drugging αPDL1 antibody and IL-12 cytokine

Thus far, we have demonstrated feasibility, safety and robust efficacy of engineering CAR T cells with PDL1-IL12 fusion proteins against solid tumors. We have also demonstrated the therapeutic benefits of engineering CAR T cells with αPDL1-IL12 versus αPDL1^mut^-IL12, supporting our initial hypothesis that sequestering the cytokine in PDL1^+^ tumors is optimal in reducing potential toxicities and enhancing therapeutic activity of IL-12. It is unclear, however, whether engineering αPDL1 and IL-12 in CAR T cells is beneficial over treatment with these modalities using biologics. To that end, we compared CAR T cells engineered with αPDL1-IL12 to CAR T cells in addition to αPDL1 antibody and IL-12 cytokine injections in the syngeneic ovarian cancer model. To mitigate the significant body weight loss that we observed with IL-12 in our previous studies and to focus more heavily on impact of systemic inflammation following therapy ^29^, we reduced the IL-12 dose when injected either alone or in combination with αPDL1. Mice were treated (i.p.) with indicated NT-CAR or TAG72-CAR T cells alone, or TAG72-CAR T engineered with αPDL1, αPDL1^mut^-IL12, or αPDL1-IL12 fusions, or TAG72-CAR co-treated with either Avelumab (which is capable of binding both human and mouse PDL1) ^57^, and/or soluble IL-12 (sIL12) (**Figure 5a**). Tumors were monitored by non-invasive imaging for 7 days, showing a trend of the most potent anti-tumor activity with TAG72-CAR/αPDL1-IL12 T cells, followed by TAG72-CAR/αPDL1^mut^-IL12 T cells and TAG72-CAR T cells combined with Avelumab and sIL12 (**Figure 5b**). Interestingly, PDL1 blockade alone, either engineered in CAR T cells or CAR T cells in combination with Avelumab, showed little to no benefit over CAR T cells alone, but showed synergy in combination with IL-12.

**Figure 5:**
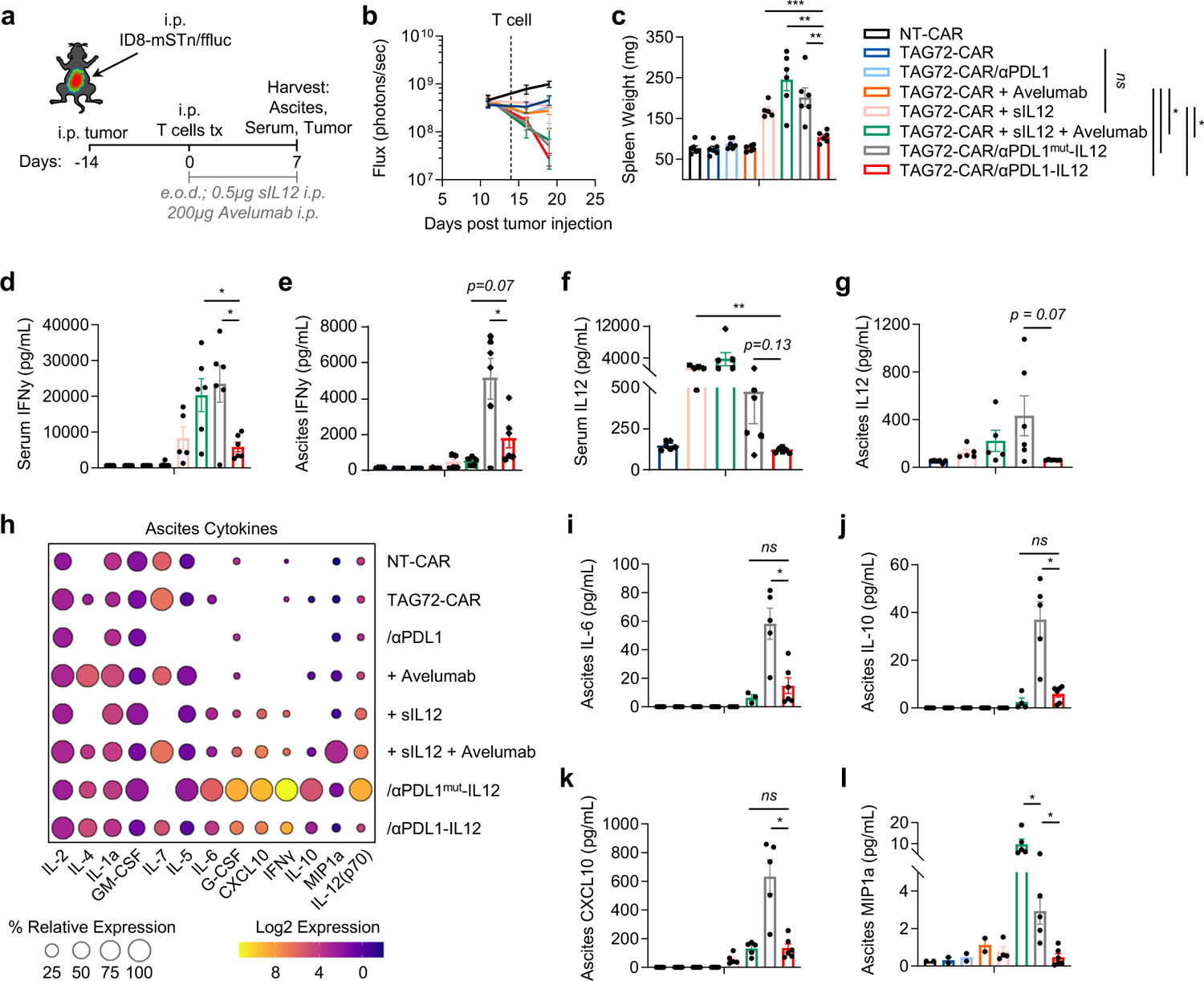
Improved safety of CAR T cells engineered with αPDL1-IL12 compared with drugging αPDL1 antibody and IL-12 cytokine. **(a)** Illustration of intraperitoneal (i.p.) ID8-mSTn/ffluc tumor model engraftment, treatment, and harvest timepoints. Mice were treated i.p. with 5.0×10^6^ NT-CAR T cells, TAG72-CAR T cells alone, or TAG72-CAR T cells engineered with αPDL1, αPDL1^mut^-IL12, αPDL1-IL12, or TAG72-CAR T cells plus exogenous treatment with soluble IL12 (+ sIL12; 0.5 µg every other day, i.p. x 3 doses), exogenous αPDL1 antibody (+ Avelumab; 200 µg every other day, i.p. x 3 doses), or in combination triple treatment (CAR + sIL12 + Avelumab) at same doses described starting on day of T cell injection. **(b)** Average tumor flux in mice treated as indicated from engraftment to just prior to euthanasia and harvests (n = 6 per treatment group). p-values for flux comparisons indicated are presented on edge of treatment legend and represent final timepoint flux prior to euthanasia. **(c)** Average spleen weights (mg) in mice harvested on day 7 post T cell treatment. IFNγ (pg/mL) by ELISA in **(d)** serum and **(e)** peritoneal ascites collected at day 7 post T cell injection. ELISA quantification of IL12 (pg/mL) in **(f)** blood serum and **(g)** peritoneal ascites collected at day 7 post T cell treatment. **(h)** Balloon plot quantifying absolute Log2 expression of mouse cytokine, and percent expression, relative to all treatment groups, measured in peritoneal ascites collected at 7 days post T cell injection in indicated treatment groups. Bar graphs quantifying individual cytokines and factors (pg/mL) in peritoneal ascites: **(i)** IL-6, **(j)** IL-10, **(k)** CXCL10, **(l)** and MIP1a. n = 5-6 mice per treatment group. All data presented as mean ± SEM. Unless otherwise indicated, p-values for pairwise comparisons were generated using an un-paired two-tailed Student’s t test with assumption of unequal variance where * p<0.05, ** p<0.01, *** p<0.001, **** p<0.0001.

Mice were subsequently euthanized at day 7 post-treatment and assessed for systemic inflammatory responses. Mice treated with TAG72-CAR T cells in combination with sIL12, sIL12 + Avelumab, and engineered TAG72-CAR/αPDL1^mut^-IL12 had significant increases in spleen weight relative to TAG72-CAR/αPDL1-IL12 (**Figure 5c**). Serum and peritoneal ascites fluid was assessed by ELISA and multiplex cytokine analysis. Treatment with TAG72-CAR/αPDL1^mut^-IL12 T cells showed significantly higher levels of serum and ascites IFNγ relative to TAG72-CAR/αPDL1-IL12 (**Figure 5d and e**) and correlated with IL12 levels (**Figure 5f and g**). Remarkably, while TAG72-CAR T cells in combination with Avelumab and sIL-12 also showed elevated serum levels of IFNy and IL-12, these cytokines were dampened in the ascites fluid compared with TAG72-CAR/αPDL1^mut^-IL12 T cell treatment. TAG72-CAR/αPDL1^mut^-IL12 T cell treatment showed a marked increase in locoregional levels of multiple factors associated with chronic or acute inflammation (**Figure 5h**) including IL-10, IL-6, CXCL10, and in triple treatment with sIL12, an increase of MIP1a, all known to be associated with cytokine release syndrome (**Figure 5i-l**). In contrast, TAG72-CAR T cells engineered with αPDL1-IL12 fusion resulted in potent anti-tumor responses and reduced systemic and locoregional cytokine levels, thereby restricting systemic inflammatory responses compared with engineering with αPDL1^mut^-IL12 or by drugging αPDL1 and IL-12. These data further support our CAR T cell engineering strategy for the safe and effective treatment of solid tumors.

## Discussion

Our study addresses a significant challenge in engineering safe and effective combinations of cytokine modification, immune checkpoint inhibition, and CAR T cell therapy for solid tumors. Our initial aim was to identify whether combining immune checkpoint inhibition of PDL1 signaling with various immune modulating cytokines (IL-12, IL-15, or TGFβ^trap^) is synergistic when engineered into solid tumor targeting CAR T cells. Using multiple *in vivo* models, we show that our CAR T cells delivering an αPDL1-IL12 fusion allows for sequestering IL-12 in PDL1+ tumors which improves safety, efficacy, and TME landscape changes. Our studies also highlight the synergistic anti-tumor activity of combining PDL1 blockade and IL-12, compared to IL-15 or TGFβ^trap^. Further, we observed an enhanced safety profile compared to CAR T cells combined with administration of a PDL1 blocking antibody and IL-12 cytokine.

Research exploiting combinations of immune checkpoint inhibitors (ICI) with cytokine modification (e.g., IL-12, IL-15, IL-18) as single agents or fusion molecules have been recently reported. As a therapeutic, αPDL1-IL15 fusions have shown mixed results in syngeneic mouse models *in vivo* ^15,27,31,32,48,58,59^, and suggests perhaps other immune checkpoints such as αPD1 or αCTLA4 may work more favorably in concert with IL-15, specifically to aid in engaging T cell or NK cell effector function and immune memory ^58^. With regards to TGFβ signaling inhibition, preclinical data also supports engaging tumor PDL1 in combination with TGFβ-“trapping” to alleviate immune suppression and boost immune and cell-based therapeutics ^16,37,60,61^. However, the αPDL1-TGFβ approach to date has faced safety challenges in phase 3 trials ^19,62^. These latter approaches, when engineered in CAR T cells, showed suboptimal safety profiles and anti-tumor activity in our models compared with αPDL1-IL12.

While effective in preclinical models as druggable agents, one major drawback of cytokine/ICI fusions is the undefined pharmacokinetics, biodistribution, and requirement of repeated administration. This also raises concerns over tumor specificity despite attempting to anchor to checkpoints, or to non-immune tumor-specific targets including cell-surface vimentin, fibronectin, or collagen binding domain ^14,18,63^. To improve tumor-specificity, Jones et al. recently described *ex vivo* loading of IL-12 anchored to CD45 expressed on antigen specific T cells prior to infusion ^33^. Their results are most closely related to our approach in that anchoring IL-12 to tumor-targeting T cells can improve anti-tumor responses, improve safety, and elicit modifications of the TME.

Our approach uses a single dose of engineered CAR T cells as a tumor-guided cellular delivery vehicle for αPDL1-IL12 fusions, enabling optimal local tumoral effects and potential for eliminating the need for repeated administration.

Our previous work established that CAR T cells engineered with membrane-bound IL-12 (mbIL12) stimulate greater IFNγ production and sequestered effects *in vivo* ^29^. While productive IFNγ signaling is crucial for effective tumor responses as well as MHC-I changes in the TME facilitated by CAR T cells and IL-12 signaling, it also can induce compensatory PDL1 expression ^6,20^. Recent findings suggest that improved immune function in immunologically “cold” cancers, such as ovarian cancer, can be achieved when combining ICIs like αPD1 and αCTLA4 with IL-12 ^64^. These studies show that these combinations drive much greater amounts of IFNγ which benefits immune function and tumor targeting via upregulation of MHC molecules. Our unique engineered fusion approach also showed enhanced IFNγ production, T cell-dendritic cell crosstalk, immune effector status, thereby presenting an opportunity to enhance the therapeutic index of ICIs in combination with CAR T cells. Most impressively, our data highlights the synergistic activity of IL12 and PDL1 blockade combined with CAR T cell therapy, as evidenced by studies combining exogenous IL12 and PDL1 blockade as well as our engineered αPDL1-IL12 fusion. And while we aimed to consolidate our treatment into a single administration, one could envision combining our mbIL12-engineered CAR T cells with PDL1 blockade, which warrants further investigation as well as in comparison to the current fusion platform.

In addition to safety and efficacy with systemic administration of CAR T cells engineered with αPDL1-IL12 fusions, we showed utility in both systemic and locoregional delivery, suggesting applicability in other regional malignancies including brain and pleural cancers ^65^. In addition, we observed that intraperitoneal delivery of unrestricted sIL12 (with or without Avelumab) with CAR T cells led to greater systemic IL-12 accumulation, while fusion-engineered CAR T cells reduced systemic IL-12 leakage and lowered systemic inflammatory effects. To further improve tumor specificity of IL-12, alternative approaches may include structural attenuation of cytokines like IL-12 ^17,49^, localized proteasomal masked cytokines ^21,34^, or logic-gating strategies for regulatable activity outside the TME ^66^. While PDL1 blockade in the presence of IL-12 promoted greater anti-tumor effects, we also observed a correlation of higher intra-tumoral CD8+ T cells with greater amounts of immune checkpoint VISTA in mice treated with TAG72-CAR/αPDL1-IL12 T cells. These data may suggest targeting other compensatory immune checkpoint pathways, like VISTA, may present a new combinatorial strategy to blunt T cell exhaustion in the TME following αPDL1-IL12-engineered CAR T cell therapy.

Overall, our study highlights a clinically translatable platform for CAR T cell engineering with localized delivery of immune modulatory αPDL1-IL12 bifunctional fusions. This approach has broad applicability to multiple solid tumors and cellular delivery holds potential across not only CAR T cells, but CAR-engineered NK cells, macrophages, and non-conventional T cells, in disease settings that warrant IC blockade and cytokine delivery to improve therapeutic responses^67–69^.

## Acknowledgements

Research reported in this publication was supported by a Prostate Cancer Foundation Tactical Award (2022TACT3825, PIs: CHJ and SJP) and the National Cancer Institute of the National Institutes of Health under grant numbers P30CA033572 and P30CA014089.

