## Supplemental Figures for "Solid tumor CAR T cells engineered with fusion proteins targeting PDL1 for localized IL-12 delivery"

Figure S1

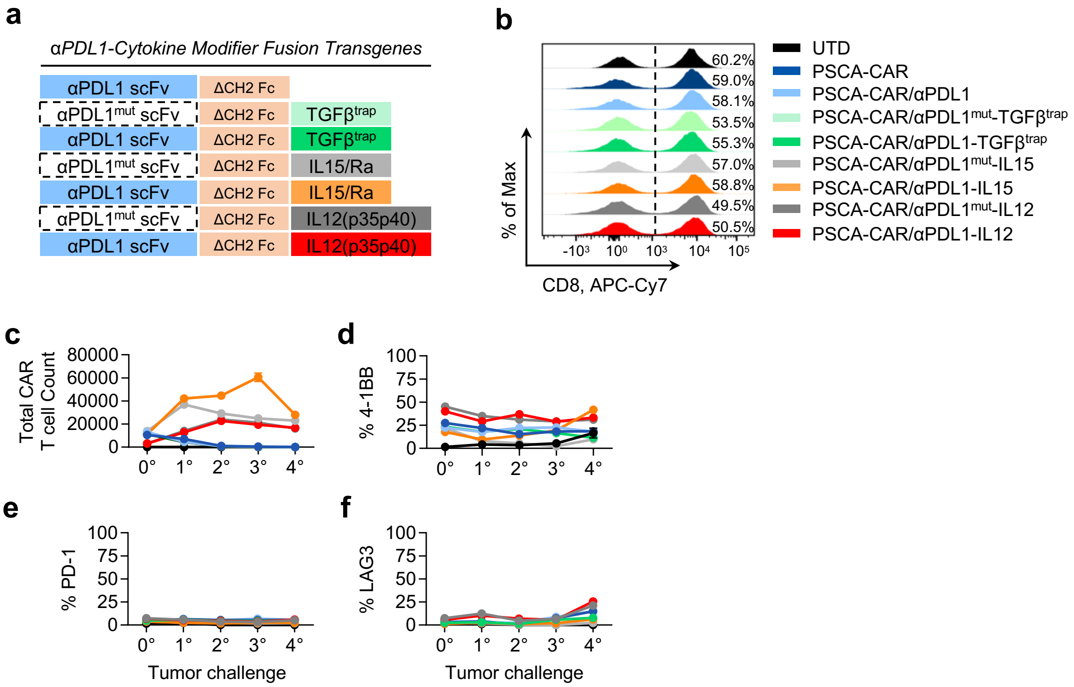

a
αPDL1-Cytokine Modifier Fusion Transgenes
αPDL1 scFv
ΔCH2 Fc
αPDL1mut scFv
TGFβtrap
IL15/Ra
IL12(p35p40)
b
% of Max
CD8, APC-Cy7
60.2%
59.0%
58.1%
53.5%
55.3%
57.0%
58.8%
49.5%
50.5%
UTD
PSCA-CAR
PSCA-CAR/αPDL1
PSCA-CAR/αPDL1mut-TGFβtrap
PSCA-CAR/αPDL1-TGFβtrap
PSCA-CAR/αPDL1mut-IL15
PSCA-CAR/αPDL1-IL15
PSCA-CAR/αPDL1mut-IL12
PSCA-CAR/αPDL1-IL12
c
Total CAR T cell Count
d
% 4-1BB
e
% PD-1
f
% LAG3
Tumor challenge

**Figure S1: (a)** Construct designs used for engineered bifunctional fusion proteins with a cytokine modifier (TGFβ$^{trap}$, IL15, or IL12) linked to an αPDL1 targeting scFv. **(b)** Flow cytometry histograms of CD8+ percentage on indicated untransduced (UTD) or PSCA-CAR conditions. Flow cytometry analysis of percentage **(c)** 4-1BB+, **(d)** PD-1+, and **(e)** LAG3+ expression on T cells, and **(f)** total CAR+ T cell count per tumor re-challenge. Data are presented as mean ± SEM. Unless otherwise indicated, p-values for pairwise comparisons were generated using an unpaired two-tailed Student's t-test with assumption of unequal variance where * $p<0.05$, ** $p<0.01$, *** $p<0.001$, **** $p<0.0001$.

Figure S2

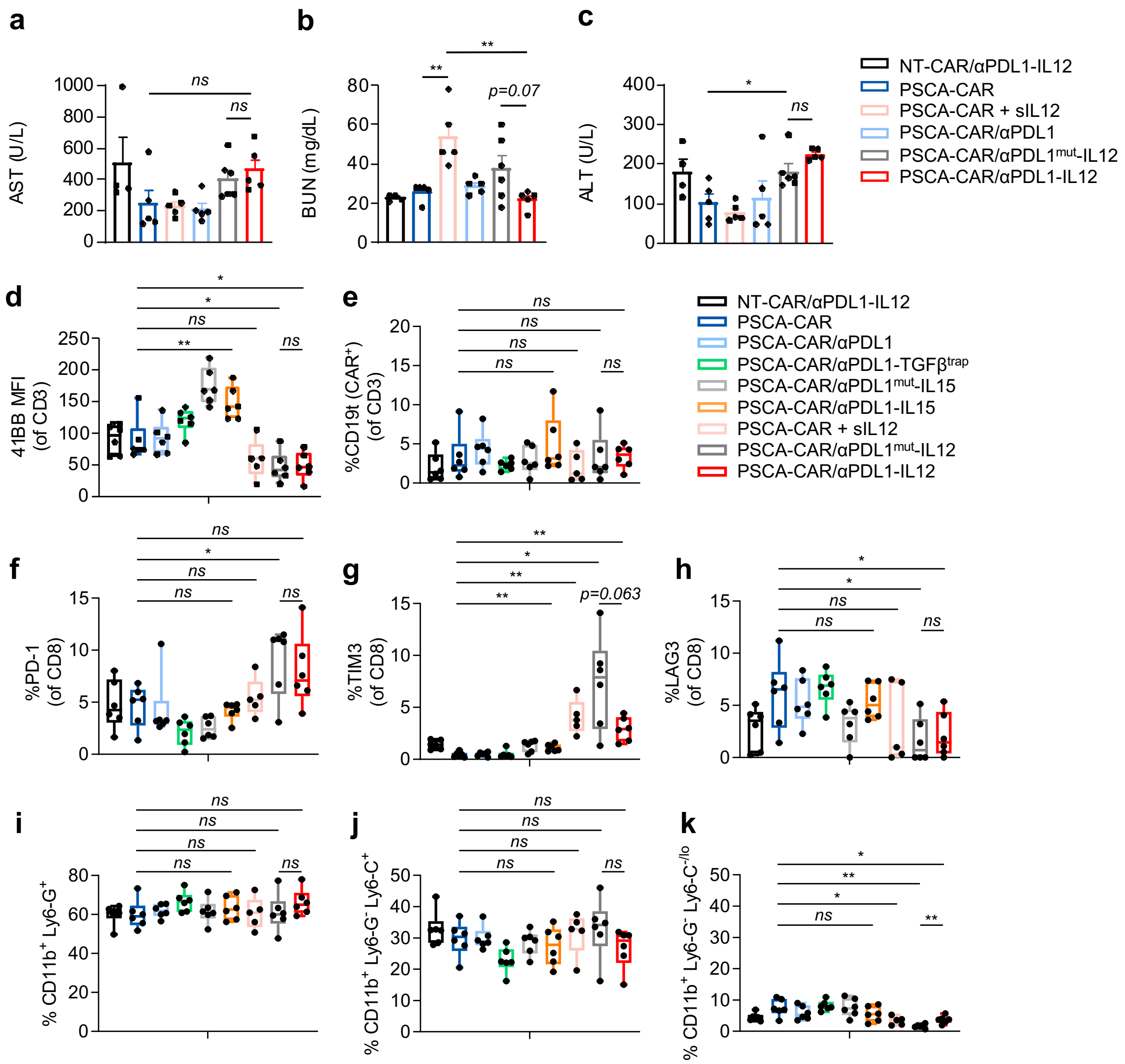

a
AST (U/L)
ns
ns
b
BUN (mg/dL)
**
**
p=0.07
c
ALT (U/L)
*
ns
NT-CAR/αPDL1-IL12
PSCA-CAR
PSCA-CAR + sIL12
PSCA-CAR/αPDL1
PSCA-CAR/αPDL1mut-IL12
PSCA-CAR/αPDL1-IL12
d
41BB MFI (of CD3)
e
%CD19t (CAR+) (of CD3)
NT-CAR/αPDL1-IL12
PSCA-CAR
PSCA-CAR/αPDL1
PSCA-CAR/αPDL1-TGFβtrap
PSCA-CAR/αPDL1mut-IL15
PSCA-CAR/αPDL1-IL15
PSCA-CAR + sIL12
PSCA-CAR/αPDL1mut-IL12
PSCA-CAR/αPDL1-IL12
f
%PD-1 (of CD8)
g
%TIM3 (of CD8)
p=0.063
h
%LAG3 (of CD8)
i
% CD11b+ Ly6-G+
j
% CD11b+ Ly6-G- Ly6-C+
k
% CD11b+ Ly6-G- Ly6-C-/lo

**Figure S2: (a-c)** Quantification of AST (U/L) **(a)**, BUN (mg/dL) **(b)**, and ALT (U/L) **(c)** in serum from indicated treatment groups. Flow cytometry analysis of peripheral blood 6 days post T cell injection for **(d)** 41BB MFI (of CD3+ T cells), **(e)** CD19t+ (% of CAR+ T cells) **(f)** PD-1 (% of CD8+ T cells), **(g)** TIM3 (% of CD8+ T cells) **(h)** LAG3 (% of CD8+ T cells), **(i)** percentage of CD11b+Ly6G+ granulocytes, **(j)** percentage of $CD11b^{+}Ly6\text{-}G^{-}Ly6\text{-}C^{+}$ monocytes, and **(k)** percentage of $CD11b^{+}Ly6\text{-}G^{-}Ly6\text{-}C^{-/lo}$ monocytes. Unless otherwise indicated, p-values for pairwise comparisons were generated using an un-paired two-tailed Student's t test with assumption of unequal variance where * $p<0.05$, ** $p<0.01$, *** $p<0.001$, **** $p<0.0001$.

### Figure S3

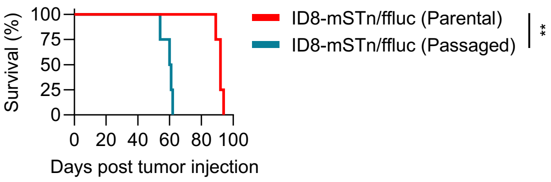
100
75
50
25
0
Survival (%)
0 20 40 60 80 100
Days post tumor injection
ID8-mSTn/ffluc (Parental)
ID8-mSTn/ffluc (Passaged)
**

**Figure S3:** Kaplan-Meier survival plot for mice engrafted i.p. with either $5.0x10^6$ parental ID8-mSTn/ffluc or aggressive passage ID8-mSTn/ffluc. n ≥ 5 mice per group, p-value using a Log-rank (Mantel-Cox) test.

Figure S4

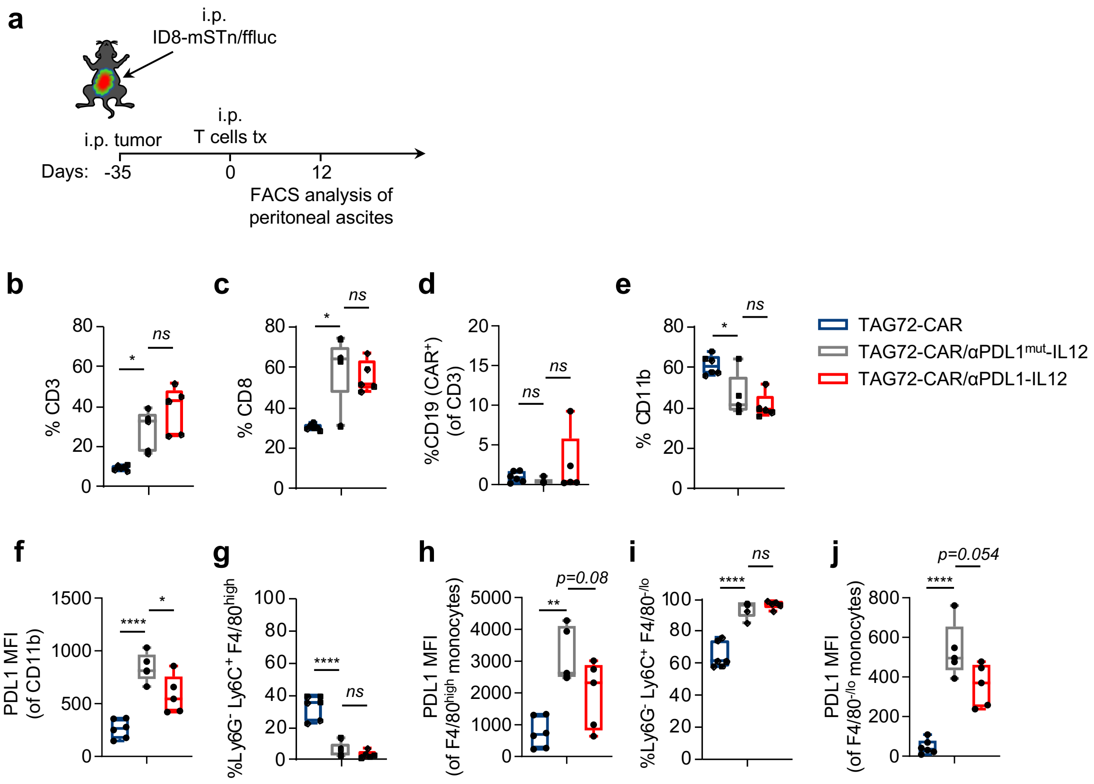
a
i.p.
ID8-mSTn/ffluc
i.p. tumor
i.p.
T cells tx
Days: -35
0
12
FACS analysis of
peritoneal ascites
b
% CD3
c
% CD8
d
%CD19 (CAR+) (of CD3)
e
% CD11b
TAG72-CAR
TAG72-CAR/αPDL1mut-IL12
TAG72-CAR/αPDL1-IL12
f
PDL1 MFI (of CD11b)
g
%Ly6G- Ly6C+ F4/80high
h
PDL1 MFI (of F4/80high monocytes)
p=0.08
i
%Ly6G- Ly6C+ F4/80-/lo
j
PDL1 MFI (of F4/80-/lo monocytes)
p=0.054
ns
*
**
****

**Figure S4: (a)** Illustration of intraperitoneal (i.p.) ID8-mSTn/ffluc tumor model engraftment, day 35 treatment, and harvest timepoint at day 12 post T cell injection as indicated. **(b-e)** Flow cytometry analysis of peritoneal ascites collected at day 12 post T cell injection, percentage of CD3+ **(b)**, CD8+ **(c)**, CD19t+ (CAR+) of total T cells **(d)**, and CD11b+ **(e)**. **(f)** Mean fluorescence intensity (MFI) of PDL1 on total CD11b+ myeloid cells, **(g)** percentage of $CD11b^{+}Ly6\text{-}G^{-}Ly6\text{-}C^{+}F4/80^{high}$ monocytes, **(h)** MFI of PDL1 on $F4/80^{high}$ monocytes, **(i)** percentage of $CD11b^{+}Ly6\text{-}G^{-}Ly6\text{-}C^{+}F4/80^{-lo}$ monocytes, and **(j)** MFI of PDL1 on $F4/80^{-/lo}$ monocytes. n ≤ 6 mice per group. Unless otherwise indicated, p-values for pairwise comparisons were generated using an un-paired two-tailed Student's t test with assumption of unequal variance where * $p<0.05$, ** $p<0.01$, *** $p<0.001$, **** $p<0.0001$.

### Figure S5

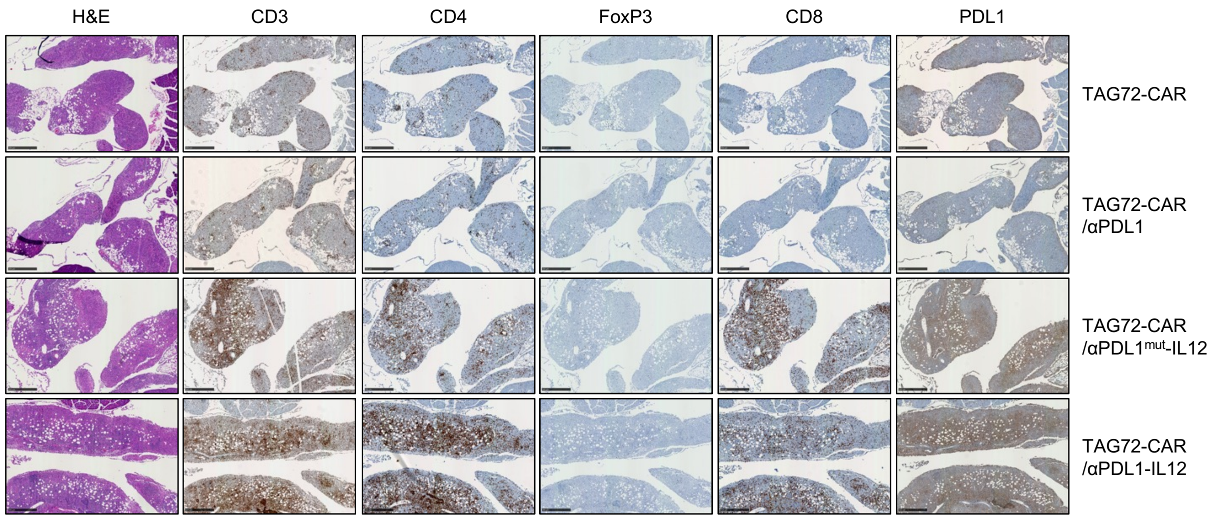
H&E
CD3
CD4
FoxP3
CD8
PDL1
TAG72-CAR
TAG72-CAR
/αPDL1
TAG72-CAR
/αPDL1mut-IL12
TAG72-CAR
/αPDL1-IL12

**Figure S5:** Representative H&E staining and immunohistochemistry of CD3, CD4, FoxP3, CD8, and PDL1 proteins on tumor tissues from ID8-mSTn/ffluc engrafted mice, collected day 7 after T cell injection in indicated treatment groups. Scale bars = 300 μm.

Figure S6

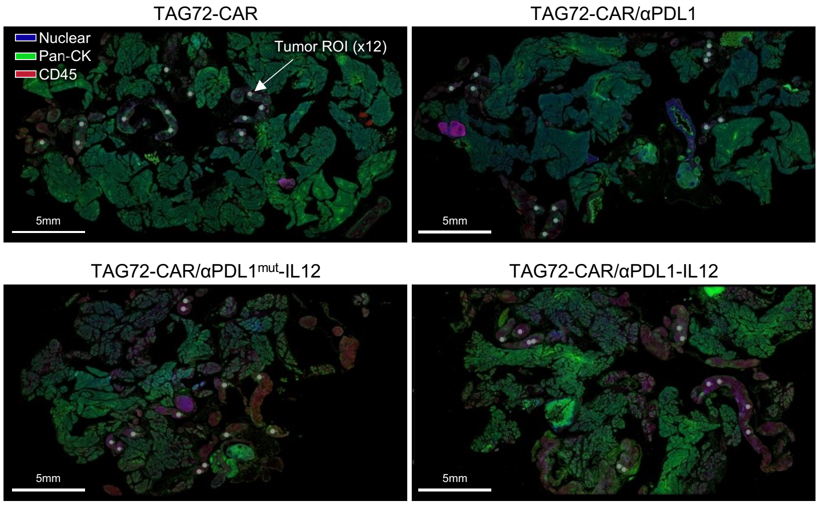
TAG72-CAR
TAG72-CAR/αPDL1
Nuclear
Pan-CK
CD45
Tumor ROI (x12)
5mm
5mm
TAG72-CAR/αPDL1^mut-IL12
TAG72-CAR/αPDL1-IL12
5mm
5mm

**Figure S6:** Total tissue Nanostring GeoMx® captured immunofluorescence images and ROI's (white dots; n = 12) from each tumor location harvested at day 7 post T cell injection from TAG72-CAR, TAG72-CAR/αPDL1, TAG72-CAR/αPDL1$^{mut}$-IL12, and TAG72-CAR/αPDL1-IL12 treatment groups. Nuclear stain (blue), Pan-CK (green), CD45 (red). Scale bars = 5 mm.
